# *Wolbachia* affect behavior and possibly reproductive compatibility but not thermoresistance, fecundity, and morphology in a novel transinfected host, *Drosophila nigrosparsa*

**DOI:** 10.1101/2020.01.21.913848

**Authors:** Matsapume Detcharoen, Wolfgang Arthofer, Francis M. Jiggins, Florian M. Steiner, Birgit C. Schlick-Steiner

**Author notes:** These authors contributed equally as senior authors (Florian M. Steiner, Birgit C. Schlick-Steiner). Corresponding author: Matsapume Detcharoen. Department of Ecology, Molecular Ecology Group, University of Innsbruck, Technikerstr. 25, 6020 Innsbruck, Austria.

## Abstract

*Wolbachia*, intracellular endosymbionts, are estimated to infect about half of all arthropod species. These bacteria manipulate their hosts in various ways for their maximum benefits. The rising global temperature may accelerate species migration and, thus, horizontal transfer of *Wolbachia* may occur across species previously not in contact. We transinfected and then cured the alpine fly *Drosophila nigrosparsa* with *Wolbachia* strain *w*Mel to study its effects on this species. We found low *Wolbachia* titer, possibly cytoplasmic incompatibility, and an increase in locomotion of both infected larvae and adults compared with cured ones. However, no change in fecundity, no impact on heat and cold tolerance, and no change in wing morphology were observed. Although *Wolbachia* increased locomotor activities in this species, we conclude that *D. nigrosparsa* may not benefit from the infection. Still, *D. nigrosparsa* can serve as a host for *Wolbachia* because vertical transmission is possible but may not be as high as in the native host of *w*Mel, *Drosophila melanogaster*.

## Introduction

*Wolbachia* are intracellular Alphaproteobacteria belonging to the Rickettsiales group (Hertig 1936; Lo *et al*., 2007). These bacteria are found in around 40-60% of arthropod species (Sazama *et al*., 2017; Zug & Hammerstein, 2012), including many species of *Drosophila* (Turelli *et al*., 2018), but *Wolbachia* diversity remains largely unknown (Detcharoen *et al*., 2019). *Wolbachia* are mainly maternally transmitted, but horizontal transfer has also been observed (Schuler et al., 2013; Werren *et al*., 2008). They have been dubbed as master manipulators (Werren *et al*., 2008) as they can manipulate their hosts biology and morphology. The four major phenotypes known are cytoplasmic incompatibility, feminization, male killing, and parthenogenesis (Werren *et al*., 2008). Among these effects, cytoplasmic incompatibility is the most studied (Werren *et al*., 2008). This effect occurs when *Wolbachia*-infected males mate with uninfected females and results in early embryonic death. It has been proposed that cytoplasmic incompatibility can promote host speciation by inducing reproductive barriers when the same host species hosts multiple, incompatible strains (Sinkins *et al*., 2005). *Wolbachia* have been shown to affect the morphology of their arthropod hosts, for example in wing size and shape (Dutra *et al*., 2016; Kriesner *et al*., 2016) and larva size (Dutra *et al*., 2016).

Depending on the particular host-strain interaction, host animals can also benefit from *Wolbachia* infection. *w*Mel-infected *Drosophila melanogaster* were reported to have higher fecundity, higher mating rate, and longer wings (Table 1). *Laodelphax striatellus* planthoppers infected with *w*Stri also had higher fecundity than uninfected ones (Guo *et al*., 2018). Bigger body size and longer lifespan were reported in *Callosobruchus chinensis* beetles infected with *w*BruCon, *w*BruOri, and *w*BruAus (Okayama *et al*., 2016). *Cimex lectularius* bedbugs require vitamin B provided by *Wolbachia w*Cle for development (Hosokawa *et al*., 2010). *Wolbachia* can also provide virus resistance in many species, including *D. melanogaster* infected with *w*Mel, *w*MelCS, or *w*MelPop (Chrostek *et al*., 2013; Teixeira, Ferreira, & Ashburner, 2008), and *w*Atab3 is required for proper oogenesis in the wasp *Asobara tabida* (Dedeine *et al*., 2004).

**Table 1.**
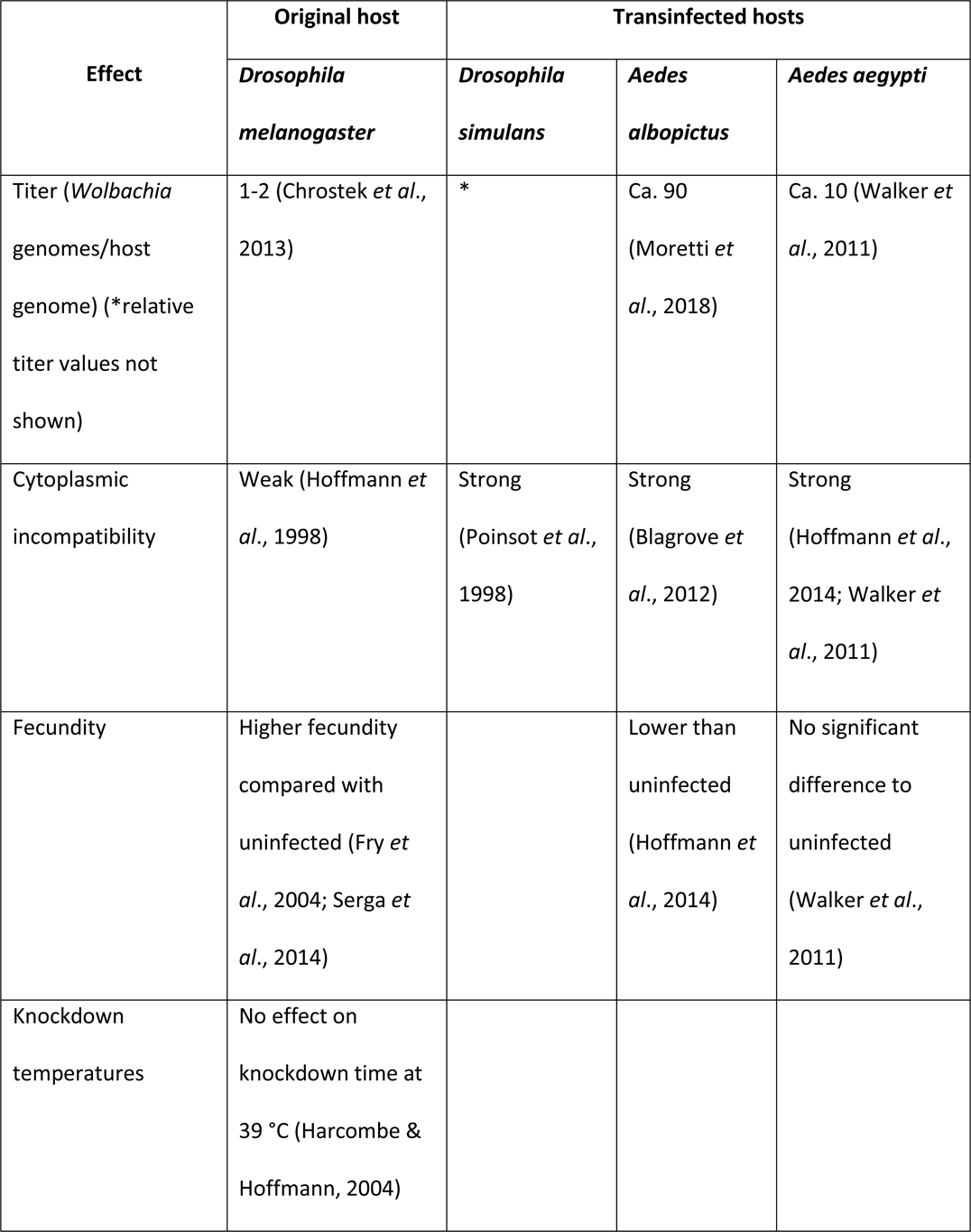
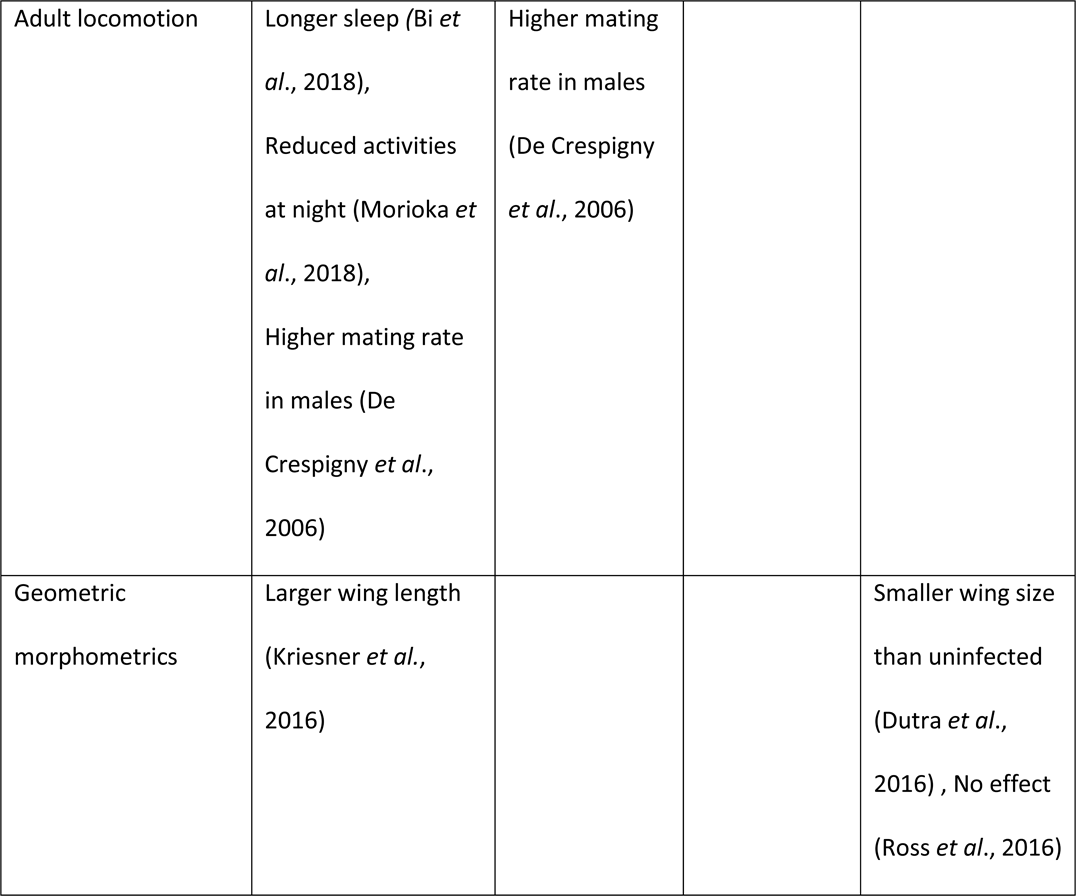
Known effects of *Wolbachia w*Mel on some infected host species.

As the global temperature has been increasing, affecting animals’ physiology and distribution (Hoffmann & Sgrò, 2011). Many animals migrate towards cooler environments (Sparks *et al*., 2005; Hickling *et al*., 2006) and reach areas previously unoccupied by these species (Dale *et al*., 2001; Parmesan & Yohe, 2003). This migration might increase the chance of animals to become infected by novel pathogens and diseases (Bebber *et al*., 2013). For example, *Wolbachia* from the European cherry fruit fly, *Rhagoletis cerasi*, has been transmitted to the invasive North American eastern cherry fruit fly, *Rhagoletis cingulata* (Schuler *et al*., 2013, 2016). In *Paratrechina longicornis*, an invasive ant species, *Wolbachia* strain *w*LonF is horizontally transmitted among populations more often than another strain, *w*LonA (Tseng *et al*., 2019).

Endosymbionts have been shown to affect hosts’ thermal biology. *Drosophila melanogaster* individuals infected with *Wolbachia w*Mel, *w*MelCS, or *w*MelPop preferred cooler temperatures compared with uninfected ones (Truitt *et al*., 2018; Arnold *et al*., 2019). Brumin *et al*., (2011) reported that *Bemisia tabaci* whiteflies infected with the endosymbiont *Rickettsia* had higher heat tolerance than uninfected ones.

*Drosophila nigrosparsa* is a species restricted to montane and alpine areas in Central and Western Europe with its main distribution around 2000 m above sea level (Bächli *et al*., 1985; Bächli *et al*., 2004). Under simulated conditions of alpine summer, these flies have around 60 days of development time from embryos to adults (Kinzner *et al*., 2016). Several life history traits and physiological limits of *D. nigrosparsa* have been studied before (Kinzner *et al*., 2018, 2016; Tratter Kinzner *et al*., 2019). *Drosophila nigrosparsa* is less fecund and relatively long living compared with other *Drosophila* species, and it is well adapted to current cold and hot temperatures (Kinzner *et al*., 2018). However, this fly species did not evolve increased heat resistance in selection experiments when maximum temperature is further increased (Kinzner *et al*., 2019). Although *Wolbachia* infect many species of *Drosophila*, all samples of *D. nigrosparsa* tested so far were uninfected (data not shown). As *D. nigrosparsa* cannot adapt to warming temperature in the recent selection experiment, they might migrate to other areas. Such migration may result in whole communities becoming mixed up, and *Wolbachia* may have the chance to encounter hitherto uninfected hosts including *D. nigrosparsa*.

Here, we focus on effects of *Wolbachia* on the new host *Drosophila nigrosparsa*. We aimed at uncovering phenotypic effects of *Wolbachia*, that is, *Wolbachia* titer fluctuation across fly ages, cytoplasmic incompatibility and fecundity, heat and cold tolerance, larval and adult locomotion, and wing geometric morphometrics. Using microinjection, three strains of *Wolbachia* commonly found in *Drosophila*, *w*Mel, *w*MelPop, and *w*MelCS, were transinfected into *D. nigrosparsa*. Subsequent generations of stably *Wolbachia*-infected, *Wolbachia*-cured, and naturally uninfected flies were characterized.

## Materials and Methods

*Drosophila nigrosparsa* naturally uninfected with *Wolbachia* was collected from Kaserstattalm in Stubai Valley, Tyrol, Austria (47.13°N, 11.30°E) in 2010 (Kinzner *et al*., 2018), and the isofemale line iso12 was established (Arthofer *et al*., 2015; Cicconardi *et al*., 2017). In this study, a subpopulation of iso12 approximately 60 generations after its establishment was used as the uninfected line nu_0 (Figure 1). *Wolbachia* status was checked using *Wolbachia* 16S (O’Neill *et al*., 1992) and wsp81F and wsp691R primers (Braig *et al*., 1998); *Cardinium* infection was checked using CLO-f1 and CLO-r1 (Gotoh *et al*., 2007) and Ch-F and Ch-R primers (Zchori-Fein & Perlman, 2004), *Spiroplasma* using primers ApDNAAF1 and ApDNAAR1 (Fukatsu *et al*., 2001) and p18-F and p18-R (Jaenike *et al*., 2010). Primers R1 and R2 (Williams *et al*., 1992) were used to test for *Rickettsia* infection.

**Figure 1.**
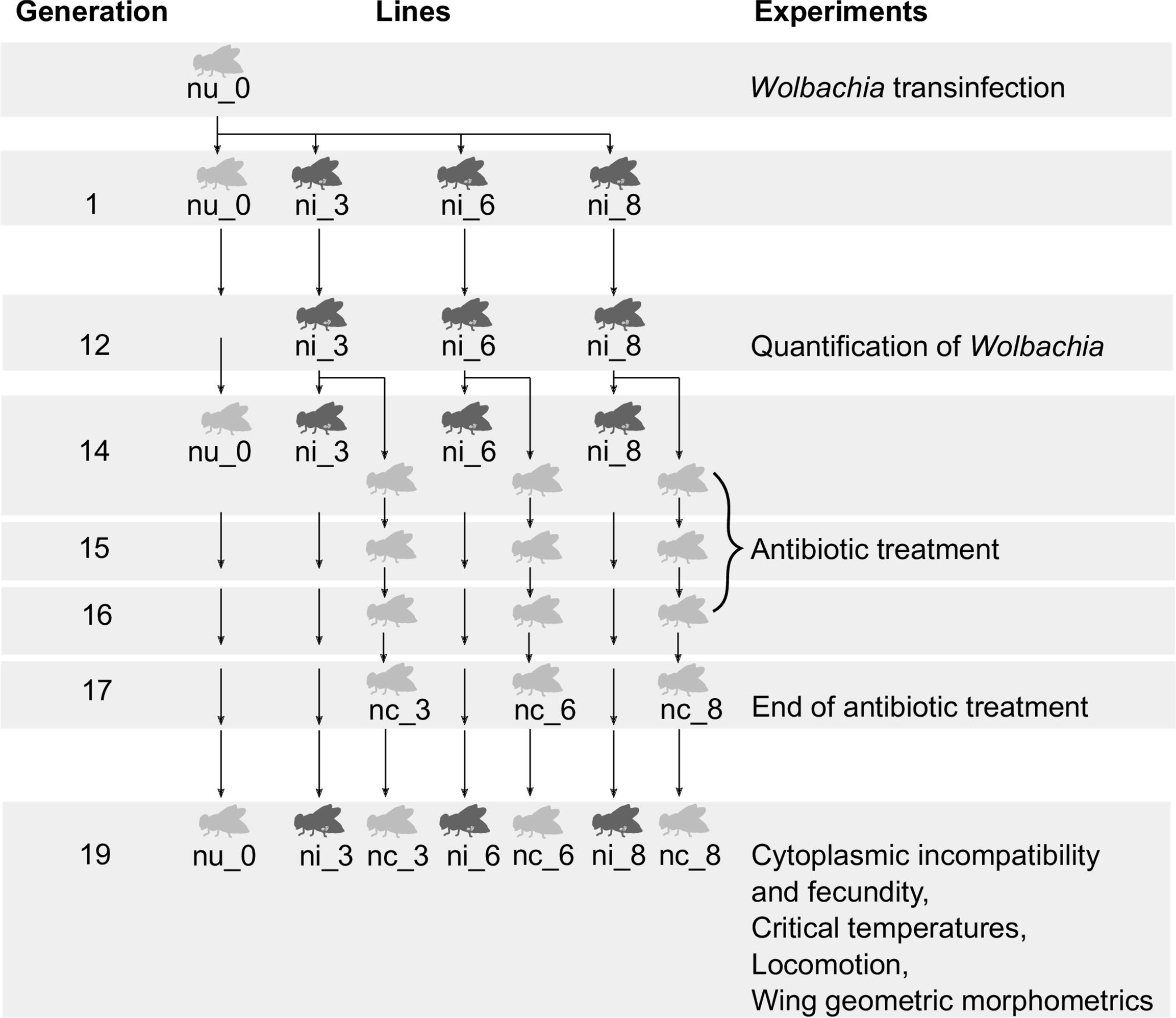
*Drosophila nigrosparsa* uninfected line nu_0 was successfully transinfected with *Wolbachia w*Mel. Generation 1 started after the establishment of three infected lines (ni_3, ni_6, and ni_8). Each fly line was kept at a census size of approximately 50 males and 50 females in every generation. Subsequent generations of uninfected and infected lines were used for the experiments.

For flies of all lines used in this study, around 50 adult males and 50 adult females were put in a mating cage modified from Kinzner *et al*. (2018) and supplied with grape-juice agar, malt food, and fresh yeast for embryo collection. Food was changed every five days. Embryos and larvae were collected and transferred to glass vials filled with malt food at a density of around 80 eggs per vial. All flies were reared at 19 °C, 70% humidity, and a 16 h:8 h light:dark cycle.

### *Wolbachia* transinfection

Cytoplasm containing a single *Wolbachia* strain from *Drosophila melanogaster* (either *w*Mel provided by Luis Teixeira, *w*MelCS, or *w*MelPop provided by Francis M. Jiggins and Julien Martinez) was injected into the posterior end of dechorionated embryos of the uninfected *D. nigrosparsa* line nu_0 at Generation 0 using a micromanipulator (M-152, Narishige, Japan) with a capillary (BF100-78-10, Sutter Instrument, USA) attached to an inverted microscope (CKX53, Olympus, Japan). Injected embryos were placed on grape-juice agar plates with fresh blobs of yeast and transferred into an incubator (MLR-352H-PE, Panasonic, Japan) for two days at 19 °C. Injected embryos developed on malt food. Each surviving female adult was mated with an uninfected male from line nu_0 to generate infected fly lines and each mating pair was kept separately in the mating cage. Three stably infected lines were generated, that is, ni_3, ni_6, and ni_8. To check for *Wolbachia* infection, five *D. nigrosparsa* females per line were randomly killed in each generation, and DNA was extracted and checked for infection by PCR using wsp81F and 691R primers.

### Quantification of *Wolbachia*

Quantitative PCR (qPCR) was used to quantify *Wolbachia* density at Generation 12. Three adult females from each stably infected line were randomly collected and checked for *Wolbachia* titer at every other day from Day 1 to Day 31 after eclosion. DNA was extracted individually from each fly using the DNeasy Blood & Tissue Kit (QIAGEN, Hilden, Germany). The relative number of *Wolbachia* cells per host cell was quantified with *Wolbachia* wsp81F and 691R primers and *D. nigrosparsa* microsatellite DN38 primers (Arthofer *et al*., 2013) as internal standard. All qPCRs were performed using Rotor-Gene SYBR Green PCR Kit (QIAGEN) with two technical replicates for each sample on a Rotor-Gene Q instrument (QIAGEN). ANOVA was used to test for titer differences among lines. All statistical analyses were done in R (R Core Team, 2018) and alpha = 0.05 was used throughout all analyses. All graphics were created using the R package ggplot2 (Wickham, 2016).

### Curing from *Wolbachia*

In Generation 14 after transinfection, two subpopulations of each stably *Wolbachia*-infected fly line were treated with tetracycline hydrochloride (lot number SLBQ2368V, Sigma-Aldrich, Germany) mixed in the malt food at final concentrations of 0.01% (Miller *et al*., 2010) or 0.05% (Schneider *et al*., 2013). After three generations of treatment, flies were transferred to normal malt food for another two generations to eliminate effects of tetracycline (Ballard & Melvin, 2007; Chatzispyrou *et al*., 2015). Three cured lines, namely nc_3, nc_6, and nc_8 were generated. Five female flies of every cured generation were randomly collected and checked for *Wolbachia* infection by PCR using the primers wsp81F and 691R.

### Cytoplasmic-incompatibility test and fecundity

The cytoplasmic-incompatibility level was assessed at Generation 19 by crossing infected, cured, and uninfected flies in all possible combinations except crosses between infected and cured flies (Table S1). Five one-day old virgin females were allowed to mate with five males of the same age from a different line in a mating cage, three cages per cross. *Drosophila nigrosparsa*, unlike *D. melanogaster* or *D. simulans*, females only lay eggs after seven days of mating and the larvae hatch two days after being laid. Thus, flies were allowed to mate for seven days. Males were removed on the eighth day, and each female was individualized into a perforated 50-ml centrifuge tube (Sarstedt, Germany) supplied with grape-juice agar, malt food, and live yeast. The number of eggs laid per female and the number of hatched larvae were counted on Day 9 and Day 14, respectively. Significance in hatching was analyzed using a generalized linear mixed model fit by maximum likelihood with a binomial error structure and logit link function implemented in the package lme4 (Bates *et al*., 2015). The number of eggs laid and the infection status of females were used as fixed effects and lines as random effect.

### Critical maximum and minimum and heat-knockdown temperatures

Critical-temperature experiments were modified from Kinzner *et al*. (2018). In Generation 19, seven-day old female flies of infected (ni_3, ni_6, and ni_8), cured (nc_3, nc_6, and nc_8), and uninfected (nu_0) lines were used. Flies were separated under carbon dioxide anesthesia two days before the experiments. On the days of experiments, flies were placed at room temperature for one hour before the experiments started and were transferred to 5-ml vials without anesthesia immediately before the experiments.

For the critical-maximum and minimum temperature assays (CTmax and CTmin, respectively), three females from the same line were transferred into a 5-ml vial, four vials per temperature. The fly-containing vials were sealed and exposed in a water bath for 5 minutes to six different temperatures from 37-39 °C for CTmax and 0.5-3.5 °C for CTmin with 0.5 °C interval. Temperatures from the thermostat reservoir (VWR, USA) and from a thermometer (Ebro TFX430, Xylem Analytics, Germany) inside the control vial were recorded with an accuracy of 0.05 °C. After 5 minutes, the vials were removed from the water bath, and the flies were checked quickly for coma by tapping the vials. Flies were discarded after each run.

For the heat-knockdown assay, three females were transferred into a 5-ml vial, four replicates per line. The vials were sealed and submerged in a transparent water bath with continuously increasing temperature from 25 °C to 39 °C at a rate of 0.47 °C min^-1^. Temperature was measured as descibed above. The number of flies in coma and the temperature inside the vials were recorded throughout the assay every 30 seconds.

The percentages of flies in coma in each vial of the CTmax and CTmin experiments were used to calculate generalized linear mixed models by maximum likelihood with a binomial error structure and logit link function of flies in coma against temperature. For heat knockdown, the temperature of each fly that was in coma was used. Analysis of covariance (ANCOVA) between infected and cured lines and *t*-test between infected and their cured lines were performed. Bonferroni correction for multiple comparisons was used.

### Locomotion

In Generation 19, 20 larvae from each infected and cured line and 31 larvae from the uninfected line were randomly collected five days after hatching. The experimental setup for assessing larval mobility was modified from Brooks *et al*., (2016). Briefly, each larva was put on 2% agarose in a 55 mm petri dish over a light pad (A4 Light Box, M.Way, China). The order of lines scored was randomized, and all larvae were recorded at the same time of the day (9-12 hours). The crawling path of each larva was recorded for three minutes using a video camera (XR155 Full HD, Sony, Japan). Total crawling distance (mm) and mean speed (mm s^-1^) were analyzed using wrMTrck plugin (Nussbaum-Krammer *et al*., 2015) implemented in Fiji (Schindelin *et al*., 2012), a version of ImageJ (Schneider *et al*., 2012) with slight modifications as described by Brooks *et al*., (2016).

The adult locomotion experiment Rapid Iterative Negative Geotaxis (RING) was modified from (Gargano *et al*., 2005). In Generation 19, fourteen-day old females from infected, cured, and uninfected lines were anesthetized with carbon dioxide for sexing and separated two days before the experiment. Ten female adults from each infected and cured line and 28 female adults from the uninfected line were used. Each female was transferred into a vial (100 x 24 x 1 mm, Scherf-Präzision Europa, Germany) and placed at room temperature an hour before the experiment. Fly-containing vials were tapped quickly so that all flies fell to the bottom, and locomotion activities (jumping and walking) were video recorded using a video camera (XR155 Full HD) for three minutes. All lines were included in each run, and the fly-containing vials were randomly placed inside the RING apparatus. All glass vials used were cleaned with heptane two days before the experiment. Stack images of the recorded videos were used for analysis using Fiji (Schindelin *et al*., 2012). The numbers of jumps and walks of each fly were counted manually.

For both larval and adult locomotion, nested ANOVAs were used to test for differences among lines within infection status. F-tests and *t*-tests were used to test between infected and their corresponding cured lines.

### Wing geometric morphometrics

To detect potential effects of *Wolbachia* infection on the morphology of *D. nigrosparsa*, two-week old female flies from infected, cured, and uninfected lines of Generation 19 were used (n = 35 for each infected and cured lines, and n = 66 for uninfected line). Both left and right wings were photographed from the upper and lower side using a Leica Z6 APO macroscope equipped with a 2.0x objective lens and a Leica MC190 HD camera connected to the Leica Application Suite version 4.0 (Leica Microsystems, Switzerland). Wing images were combined into a tps file using tpsUtil64 version 1.76 (https://life.bio.sunysb.edu/morph/soft-utility.html). Thirteen landmarks on each wing (Figure 6B) were marked manually using tpsDig2 version 2.31 (https://life.bio.sunysb.edu/morph/soft-dataacq.html).

MorphoJ version 1.06d (Klingenberg, 2011) was used to process the tps file. Images were aligned by the principal axis. Outliers were removed as detected by the cumulative distribution of the squared Mahalanobis distance. The potential imaging error between the upper and lower sides of the wings was accessed using Procrustes ANOVA. The average Procrustes coordinates of upper and lower sides of each wing and between left and right wings of each individual fly were computed. The covariance matrix was used to generate Principal Component Analyses (PCA) with 10,000 permutations.

Regression of Procrustes distance against centroid size, pooled within lines and infection status, was calculated. The residuals from the regression between Procrustes coordinates and centroid size were used for Canonical Variate Analysis (CVA) of all fly lines with 10,000 permutations. Procrustes ANOVA was performed. Asymmetry on size and shape between left and right wings between infected and cured lines was calculated as previously described (Padró *et al*., 2014). In brief, Pearson correlations were used between mean individual wing size and the difference between left and right wings for size asymmetry, and Procrustes ANOVA of wing shape was used for shape asymmetry.

## Results

*Drosophila nigrosparsa* line nu_0 was found to be not infected with *Wolbachia, Cardinium*, *Spiroplasma*, and *Rickettsia* before the start of our experiments.

### *Wolbachia* transinfection

Of all injected embryos, only embryos of *D. nigrosparsa* nu_0 injected with *Wolbachia w*Mel survived to adults. From 396 injected embryos, 145 larvae hatched, and 89 of them eclosed. Of these, 39 were females. Stable *w*Mel infection was detected after three, six, and seven generations for lines ni_3, ni_6, and ni_8, respectively.

For the other two *Wolbachia* strains, we injected 1,333 embryos with *Wolbachia* strain *w*MelPop (11 attempts) and 2,093 embryos with *w*MelCS (16 attempts). None of the embryos injected with *w*MelPop survived, and only two adult flies injected with *w*MelCS eclosed. We observed that most injected embryos died as larvae, and a few larvae injected with wMelCS died during pupation.

### Quantification of *Wolbachia*

The *Wolbachia* titer of all infected lines of Generation 12 was generally low. We observed, on average (mean ± standard error), 0.04 ± 0.01, 0.06 ± 0.01, and 0.06 ± 0.01 *Wolbachia* genomes per fly genome in the first 13 days for ni_3, ni_6, and ni_8, respectively (n = 21 per line). *Wolbachia* titer increased and reached the highest density after the second week (Figure 2). In general, line ni_8 had lower *Wolbachia* titer than lines ni_3 and ni_6.

**Figure 2.**
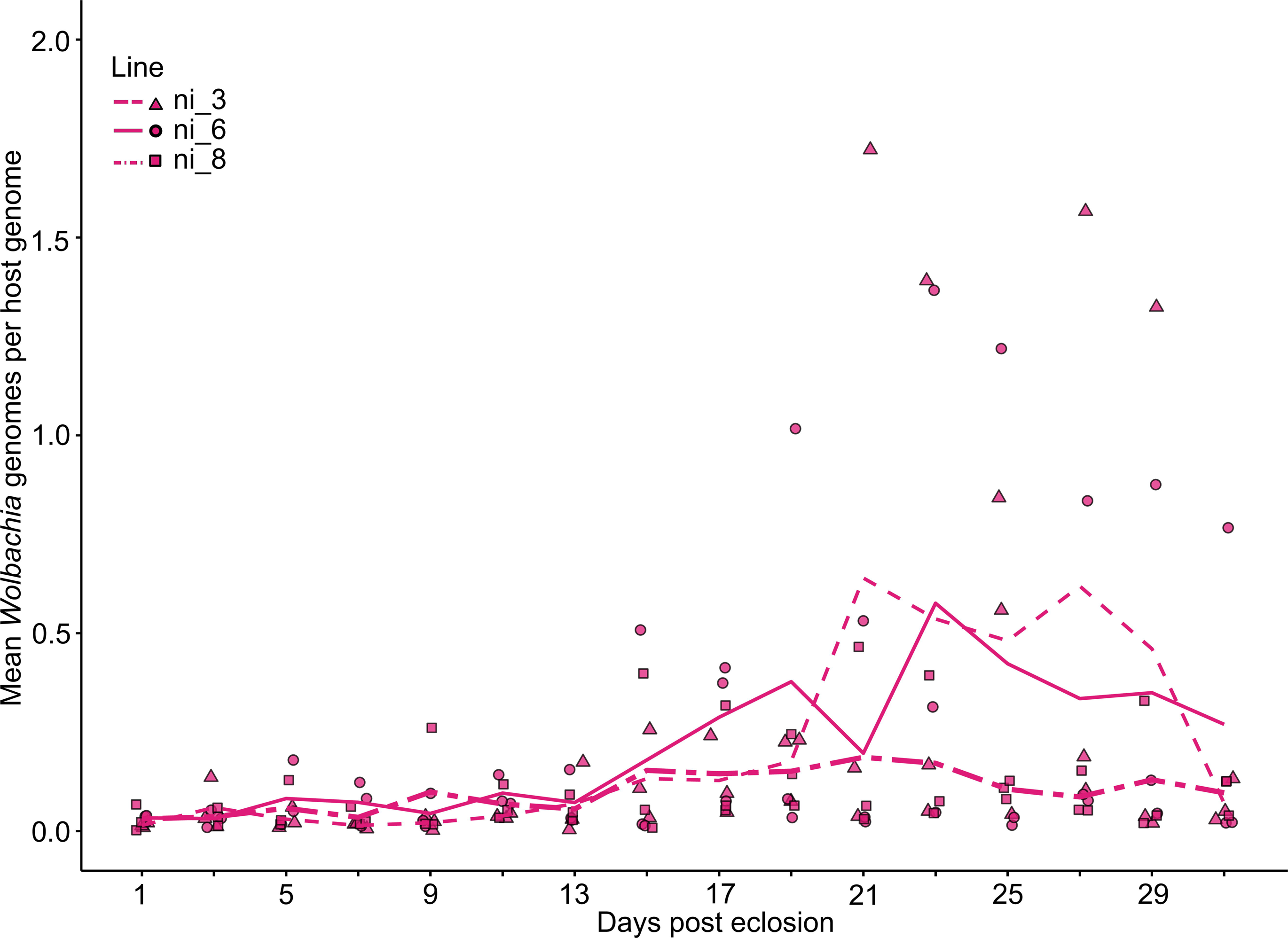
*Wolbachia* titer at different ages after eclosion of the *Wolbachia w*Mel-infected *Drosophila nigrosparsa* lines ni_3, ni_6, and ni_8 of Generation 12 quantified using qPCR. Three female flies per infected line were collected every other day. Each data point represents a biological replicate, two technical replicates were measured for each sample, and lines are mean *Wolbachia* titer. *Wolbachia* titer in all three lines varied as flies aged.

### Curing from *Wolbachia*

We did not detect *Wolbachia* with PCR during the treatment with 0.01 or 0.05% concentrations of tetracycline. However, we detected *Wolbachia* in all lines in the first generation after having stopped treating the flies with 0.01% tetracycline. *Wolbachia* were successfully removed with 0.05% tetracycline. The third generation of flies after treatment with 0.05% tetracycline was used for further experiments.

### Cytoplasmic incompatibility and fecundity

Each female laid on average (mean ± standard error) between 9.3 ± 3.5 and 15.7 ± 4.6 eggs for crosses between infected lines, 8.2 ± 2.0 and 13.0 ± 2.7 eggs between cured lines, and 12.1 ± 1.6 eggs for uninfected line (Table S2). There was no significant difference in eggs laid among lines (ANOVA, *F*_6,94_ = 0.68, *P* = 0.67) and between infection statuses of female flies (Generalized linear model, z = - 1.22, *P* = 0.22).

Crosses between infected males and females yielded similar percent hatch per cross to those between uninfected flies (mean ± standard error: 73.6 ± 6.1% and 84.7 ± 8.9%, respectively). Hatch rate dropped from 60.7 ± 5.4% in crosses of uninfected males with infected females (expected compatible cross) to just 37.7 ± 3.8% in crosses of infected males with uninfected females (expected incompatible cross) (Figure 3), but these two groups were not significantly different (Generalized linear model, z = -1.35, *P* = 0.18).

**Figure 3.**
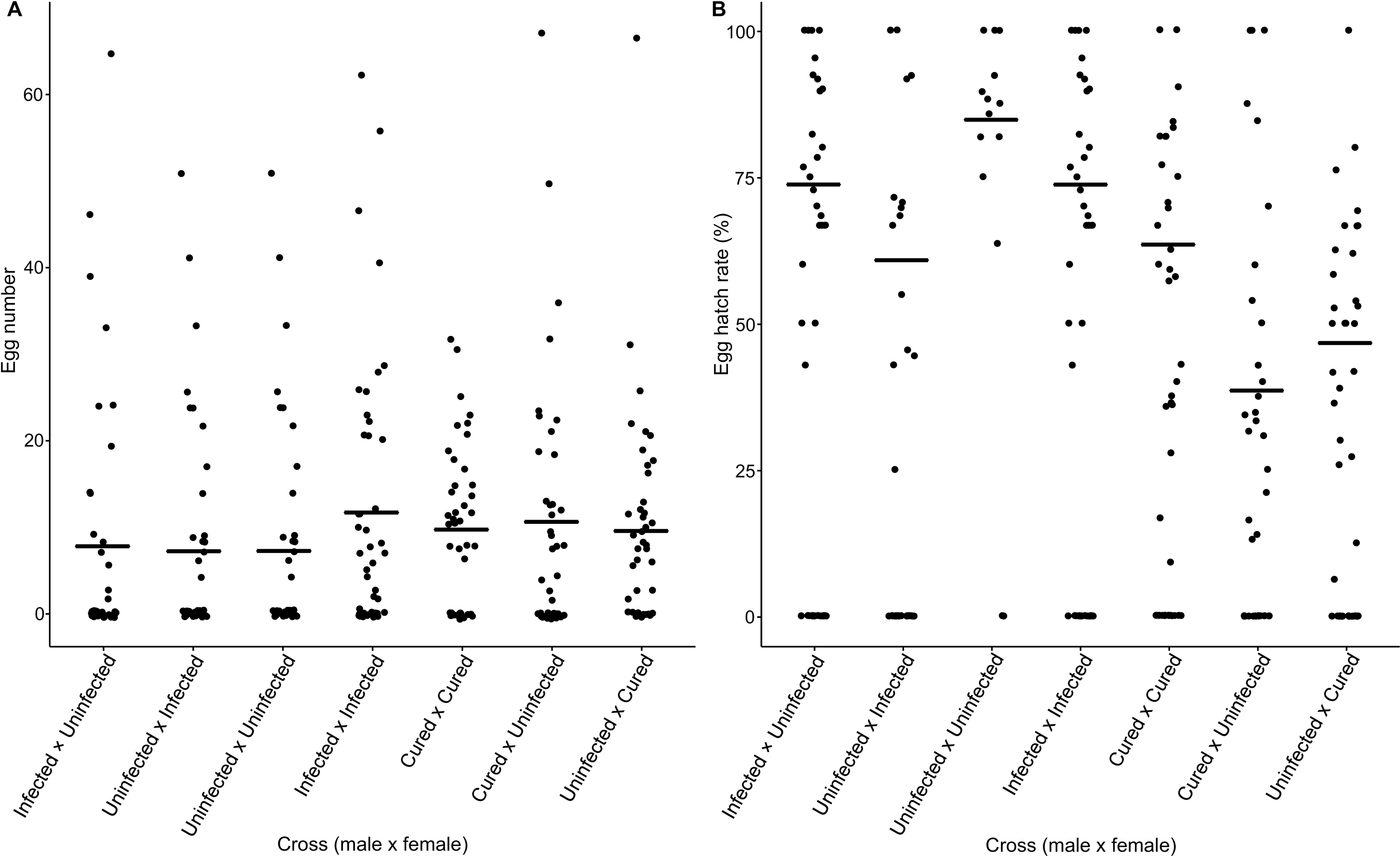
Number of eggs laid (A) and percent egg hatch of crossing between male and female of each group (B). Flies were allowed to mate for eight days, and the numbers of eggs and of hatched larvae were counted on Day 9 and Day 14, respectively. The numbers of eggs laid were not significantly different among crosses. The hatch rates were reduced in crosses of uninfected males with infected females compared with crosses of infected males with uninfected females.

### Critical-maximum and minimum and heat-knockdown temperatures

For CTmax and CTmin, generalized linear mixed models of the numbers of flies in coma against temperatures were significant in many lines (Table 2). Temperatures at which fifty percent of flies fell in coma (CT_50%_) were between 37.89 and 38.35 °C for CTmax and between 1.54 and 2.23 °C for CTmin. We did not observe any statistically significant difference between infected and cured lines in responses to temperature for CTmax (ANCOVA, *F*_1,94_ = 0.10, *P* = 0.76) nor CTmin (ANCOVA, *F*_1,94_ = 0.42, *P* = 0.52). *t*-tests between infected and corresponding cured lines were not significant (Table S2).

**Table 2.**
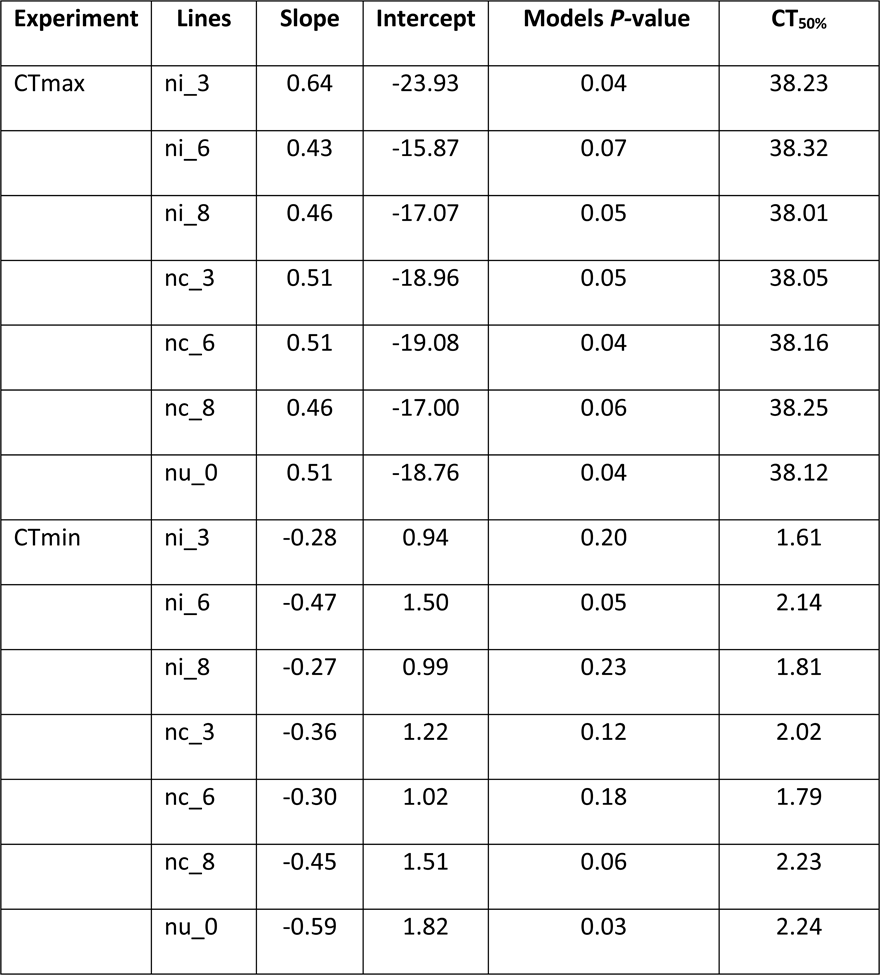
Generalize linear models of critical maximum (CTmax) and minimum (CTmin) temperatures, and temperature, at which 50% of flies fell in coma (CT_50%_).

For heat knockdown, flies fell in coma, on average, between 37.88 ± 0.98 °C from line ni_8 and 38.66 ± 0.61 °C from line nc_6 (Figure 4). Knockdown temperatures did not differ significantly between infected and cured lines (ANCOVA, *F*_1,65_ = 1.52, *P* = 0.22).

**Figure 4.**
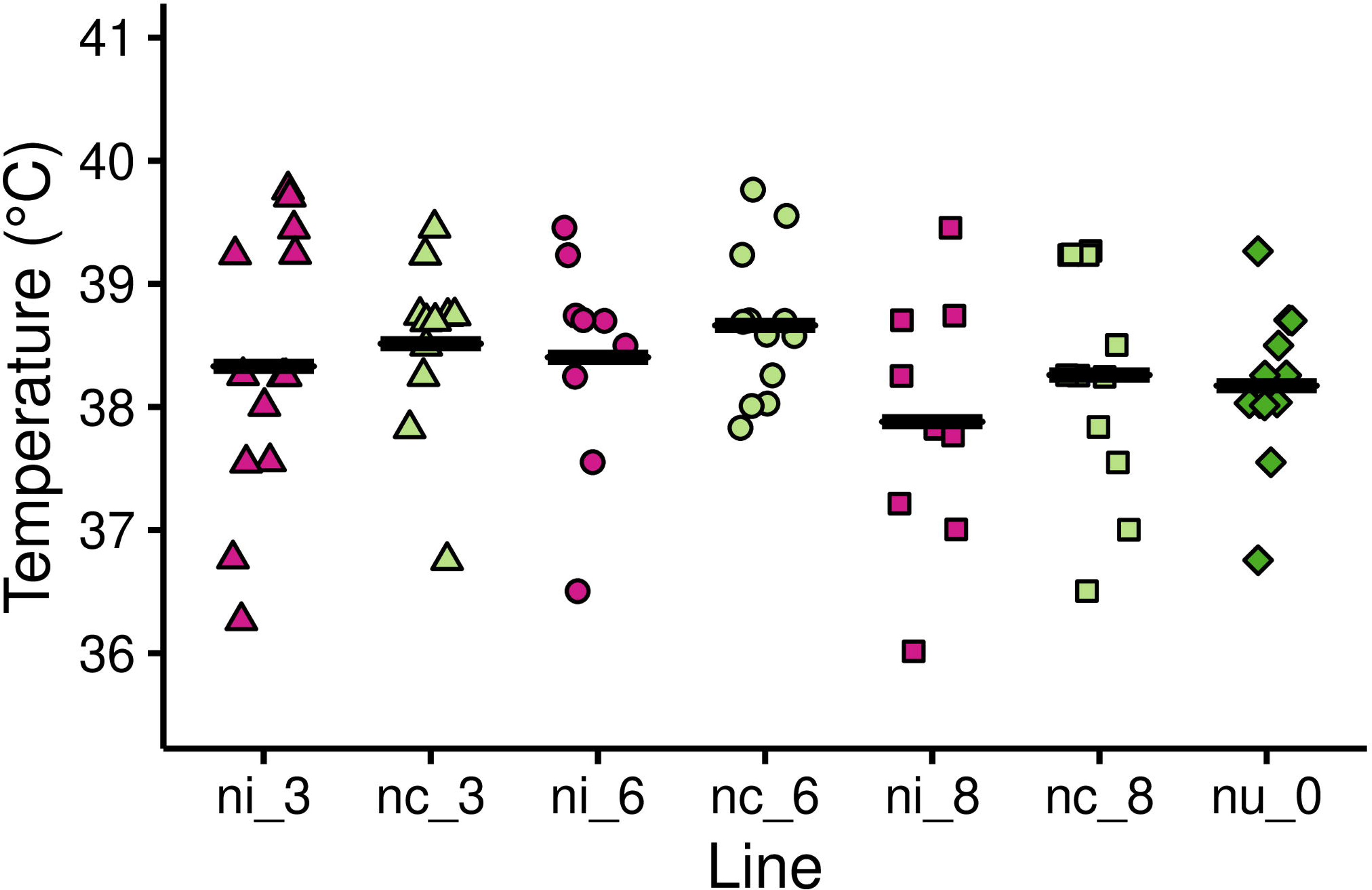
Heat knockdown temperatures of 7-day old female *Drosophila nigrosparsa* infected (ni_3, ni_6, and ni_8), cured (nc_3, nc_6, and nc_8), and uninfected (nu_0) adults. Black bars indicate mean knockdown temperatures.

In all three experiments, we found no significant different between flies of cured and uninfected lines (ANCOVA, *F*_1,62_ = 0.22, *P* = 0.88, *F*_1,62_ = 0.30, *P* = 0.58, and *F*_1,47_ = 2.49, *P* = 0.12, for CTmax, CTmin, and knockdown, respectively).

### Locomotion

For larval locomotion, infected line ni_8 had the highest mean crawling speed and the longest mean distance (Figure 5A and B, respectively). This infected line crawled significantly faster (*t*-test, *P* < 0.01) and had longer distances (*P* < 0.01) compared with its cured counterpart, nc_8. Nonetheless, mean crawling speed and crawling distances did not differ significantly between infected and cured lines (Nested ANOVA, speed, *F*_1,4_ = 0.87, *P* = 0.40; distance, *F*_1,4_ = 1.68, *P* = 0.26). In addition, we found no difference between cured and uninfected lines in both larval activities (Nested ANOVA, average speed *F*_1,2_ = 0.44, *P* = 0.58 and total length *F*_1,2_ = 1.40, *P* = 0.36).

**Figure 5.**
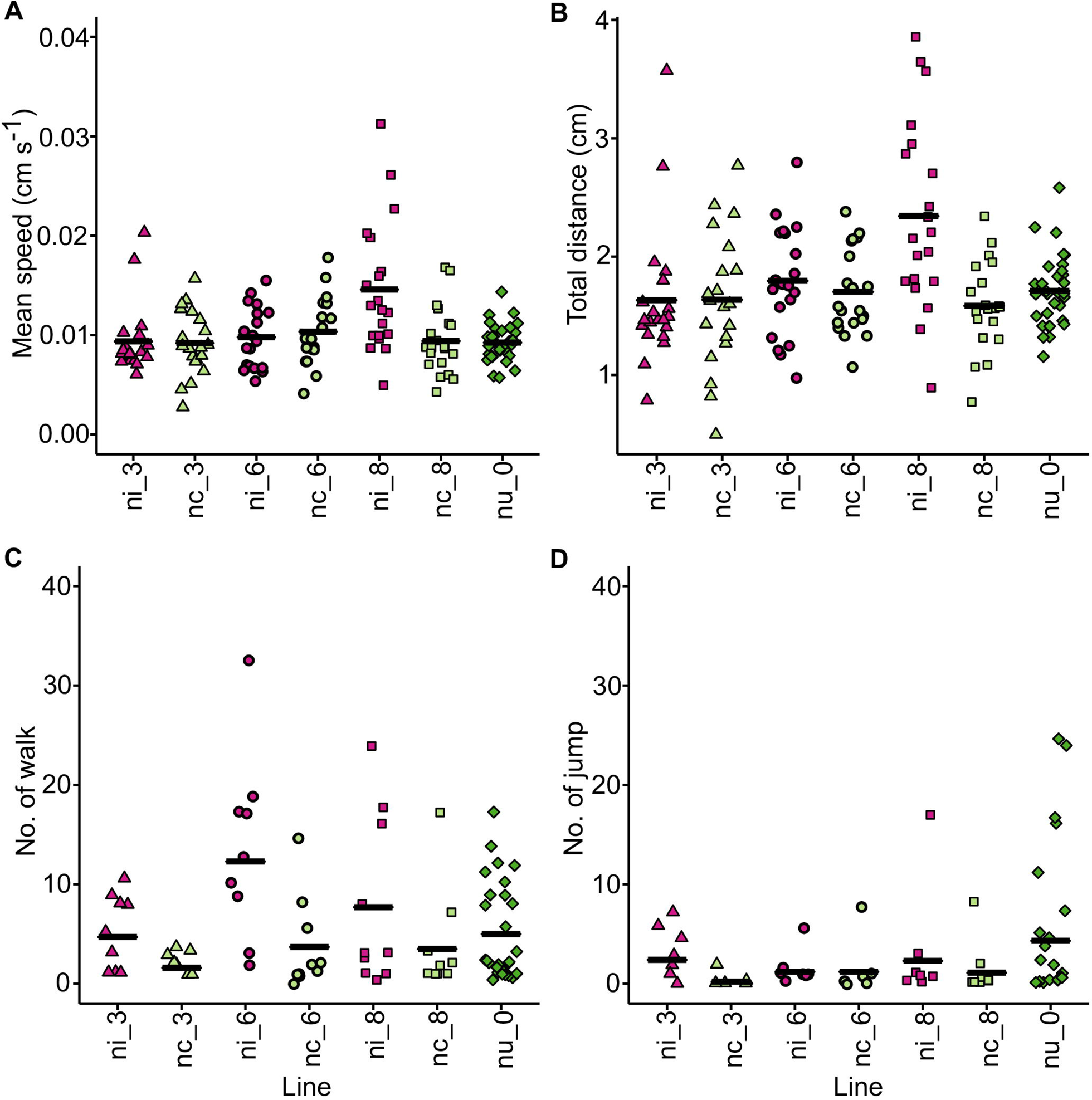
Mean speed (A) and total distance (B) of larvae crawled in three minutes (N = 10 each for infected and cured lines, 28 for uninfected line) and walk (C) and jump (D) activities of adult flies (N = 20 each for infected and cured lines, 31 for uninfected line). Plots show different Y-scales.

**Figure 6.**
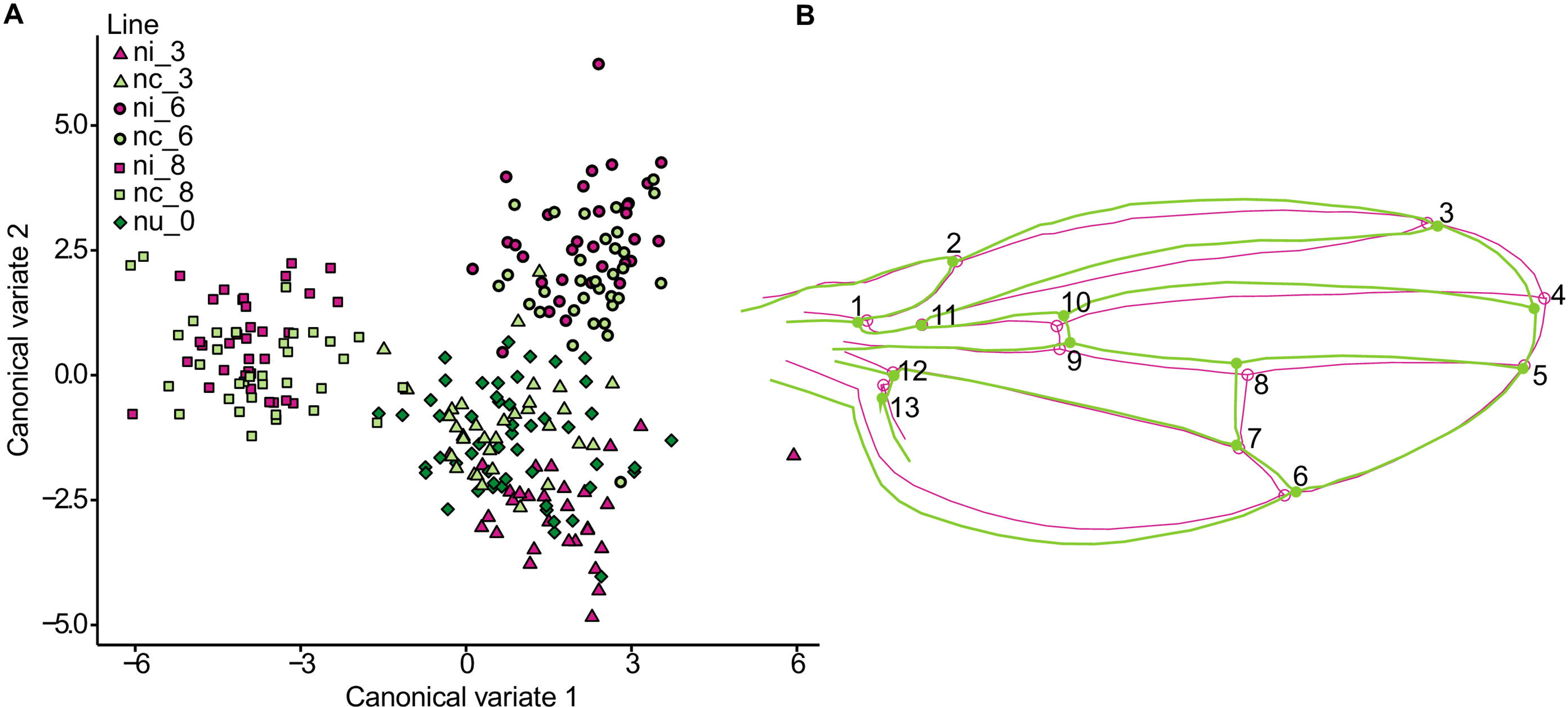
(A) Canonical variate analysis of wings of infected, cured, and uninfected lines. Each dot represents an individual fly. (B) No significant differences in average shape of all cured (green) and infected lines (pink). The differences were magnified ten times and all thirteen landmarks are shown.

In adults, infected lines had higher activities than cured lines (Figure 5C, D). Lines ni_3 and nc_3 differed significantly in both walk and jump activities (*t*-test, *P* = 0.04 and 0.03 respectively). Lines ni_6 and nc_6 differed significantly in walk activity (*t*-test, *P* = 0.03). No significant difference for walk (Nested ANOVA, *F*_1,4_ = 5.27, *P* = 0.08) nor jump activity (Nested ANOVA, *F*_1,4_ = 5.16, *P* = 0.09) between infected and cured lines was found. Comparison between cured and uninfected lines found no significant difference between cured and uninfected lines in adult walk activity (Nested ANOVA, *F*_1,2_ = 4.61, *P* = 0.17), but we found that uninfected flies had higher jump activity in uninfected than cured flies (*F*_1,2_ = 58.09, *P* = 0.02). However, between infected and uninfected lines, there was no difference in both walk and jump activities (*F*_1,2_ = 1.03, *P* = 0.42 and *F*_1,2_ = 18.11, *P* = 0.05, respectively).

### Wing geometric morphometrics

The imaging of wings can be assessed as done accurately, in that the mean squares of imaging error were very low for both centroid size and shape (2.75 and 4.54 times lower than individual by side interactions for centroid size and shape, respectively). We found significant difference among fly lines in both size and shape of the wings (Procrustes ANOVA, size, *F*_6,1_ = 456.5, *P* < 0.001; shape, *F*_132,22_ = 99.1, *P* < 0.001). CVA after removing 6.5% of total variation within lines, calculated from regression, revealed that infected and the corresponding cured lines were similar to each other (Figure 6A).

There was no difference in average shape of all cured and infected lines (Figure 6B). When comparing infected and its corresponding cured lines, we observed significant changes in centroid size and shape between ni_3 and nc_3 (size, *F*_1,1_ = 509.87, *P* = 0.03; shape, *F*_22,22_ = 104.27, *P* < 0.01) and ni_6 and nc_6 (size, *F*_1,1_ = 4815.15, *P* = 0.01; shape, *F*_22,22_ = 26.99, *P* < 0.01), and significant difference in shape between ni_8 and nc_8 (shape, *F*_22,22_ = 2.78, *P* = 0.01). Centroid size and shape of infected and cured lines differed significantly from naturally uninfected line nu_0 (Table S3 and Table S4, respectively). However, there was a small distance between groups relative to within-group variation (Mahalanobis distance = 1.20), and most flies were assigned into wrong groups (Table S5). There was no significant difference in size and shape asymmetry between left and right wings (Table S6 and Table S7, respectively).

## Discussion

Transinfection is a useful tool to investigate effects of *Wolbachia* on new host species (Hughes & Rasgon, 2014), in that it provides possibility to study a broad range of phenotypic effects on the host. We used transinfection to study effects of *Wolbachia* on *D. nigrosparsa* because this fly species may become infected by *Wolbachia* in the future by horizontal transmission upon contact with other arthropod species as a result of climate-change triggered migration. We successfully transinfected *Wolbachia w*Mel into embryos of *D. nigrosparsa* but failed to transinfect two other strains, *w*MelPop and *w*MelCS.

There are various potential reasons for the unsuccessful transinfection of *w*MelPop and *w*MelCS into *D. nigrosparsa*. Although all *Wolbachia* strains used in our study are closely related (Riegler *et al*., 2005; Woolfit *et al*., 2013), they differ in pathogenicity (van den Hurk *et al*., 2012; Woolfit *et al*., 2013). Both *w*MelPop and *w*MelCS are more virulent than *w*Mel, have higher titer inside their host, and cause early death in *Drosophila melanogaster* (Chrostek *et al*., 2013; Min & Benzer, 1997). In addition, higher pathogenicity was observed in *w*MelPop when transinfected into *Drosophila simulans* and *Aedes albopictus* compared with its native host, *D. melanogaster* (McGraw *et al*., 2001; Suh *et al*., 2009). High autophagic activity against *Wolbachia* in a novel host (Le Clec’h *et al*., 2012) might also explain our unsuccessful transinfection.

*Wolbachia* titer in their hosts depends on numerous factors. The same *Wolbachia* strain can have different titers in different host genotypes (McGraw *et al*., 2002; Lu *et al*., 2012; Early & Clark 2013) (Table 1), and, within a host, titers vary among tissues such that, for example, higher titers were observed in reproductive than in somatic tissues (Martinez *et al*., 2015; Osborne *et al*., 2012). In addition, *Wolbachia* titer might be higher if *D. nigrosparsa* was raised at a temperature cooler than 19 °C, as in our experiment, because higher *Wolbachia* density was detected in *D. melanogaster* developed at cool temperatures than those developed at warm temperatures (Moghadam *et al*., 2018).

Titer can also change with host age as observed in many arthropods including *Drosophila* spp. (McGra*w et al*., 2002; Unckless *et al*., 2009; Tortosa *et al*., 2010; Chrostek *et al*., 2013). The *Wolbachia* titer we observed (Figure 2) is likely to correlate with egg laying activity in *D. nigrosparsa*, which was reported to peak between the second and the fourth week (Kinzner *et al*., 2018). As *Wolbachia* are mainly found within host’s reproductive tissues (Frydman *et al*., 2006; Werren, 1997), the declining of *Wolbachia* titer when the flies neared completion of their fourth week could be explained by the declining of germline stem cell division with increasing individual age (Zhao *et al*., 2008).

To cure *D. nigrosparsa* from *Wolbachia*, we tried two tetracycline concentrations, 0.01 and 0.05%. High tetracycline concentration has been reported to have negative effects on hosts during the process of curing, such as fitness (Miller *et al*., 2010), and lower concentrations should therefore be preferred. However, the 0.01% concentration was too low to eliminate *Wolbachia*, which effect has likewise been reported for *Wolbachia*-infected *Drosophila paulistorum* (Miller *et al*., 2010). In addition, both *D. nigrosparsa* treated with 0.01% and 0.05% tetracycline suffered from low fecundity and low hatch rates (data not shown). We waited for another two generations before using them for our remaining experiments to recover flies from tetracycline because effects of tetracycline on mitochondrial density and metabolism can last up to two generations after treatment (Ballard & Melvin, 2007).

We noted that the recovering time of hosts after being treated with antibiotics is important. Effects of antibiotics on the hosts were eliminated within generations after treatment (Chaplinska, Gerritsma, Dini-Andreote, Falcao Salles, & Wertheim, 2016; Fry, Palmer, & Rand, 2004; H. Noda et al., 2001). However, in *Wolbachia*-infected *D. simulans*, the effects of antibiotics on productivity were eliminated five generations after treatments in two of the three lines, but these effects were not found in other traits such as egg-to-adults development time and viability (Poinsot & Mercot, 1997). To further confirm effects of antibiotics on *D. nigrosparsa*, comparisons between uninfected and uninfected flies treated with tetracycline should be further tested.

Cytoplasmic incompatibility is the most commonly observed phenotype of *Wolbachia* on their hosts (Werren *et al*., 2008). *Wolbachia w*Mel possibly induced weak cytoplasmic incompatibility in *D. nigrosparsa*, as hatchings from crosses of infected males with uninfected females (expected incompatibility) and uninfected males with infected females (expected compatibility) were reduced, although there was no difference in the number of eggs laid. Increasing the number of compatible and incompatible crosses would be needed to decide whether the lack of statistical significance in the data presented here is due to the lack of a biological effect or due to the effect being just weak; for technical reasons, additional crosses are impractical at the point of writing this manuscript. In contrast to our results, *Wolbachia w*Mel, once transinfected into other hosts, induced a high level of incompatibility, such as in *Drosophila simulans* (Poinsot, Bourtzis, Markakis, Savakis, & Merçot, 1998), in the whitefly *Bemisia tabaci* (Zhou & Li, 2016) and in the mosquito *Aedes aegypti* (Hoffmann *et al*., 2014; Walker *et al*., 2011) (Table 1).

The levels of cytoplasmic incompatibility depend on many factors. A high level of cytoplasmic incompatibility has been reported to positively correlate with high *Wolbachia* titer (Bourtzis *et al*., 1996; Noda *et al*., 2001). Young males and a high number of infected sperms also caused high level of cytoplasmic incompatibility (Clark *et al*., 2003; Reynolds & Hoffmann, 2002; Veneti *et al*., 2003). For example, Reynolds & Hoffmann, (2002) found a lower level of incompatibility when using five-day old males for crossing than using one-day old males.

The ability to adapt to elevated temperatures is an important criterion for species distribution in *Drosophila* (Kellermann *et al*., 2012). A previous study found no effect on heat knockdown temperature in *w*Mel-infected *Drosophila melanogaster* (Harcombe & Hoffmann, 2004). This finding for *Wolbachia* contrasts one for *Rickettsia*, which were reported to increase heat shock tolerance in *Bemisia tabaci* to up to 40 °C (Brumin *et al*., 2011). In *D. nigrosparsa*, a recent selection experiment on naturally uninfected flies reported that this species is unlikely to adapt to increasing temperature (Kinzner *et al*., 2019). Here, we conclude that infection with *Wolbachia w*Mel did not increase heat and cold tolerance in *D. nigrosparsa*. *Wolbachia*-infected and *Wolbachia*-free *D. nigrosparsa* responded to knockdown temperature at around 38 °C like in an earlier study of this fly species (Kinzner *et al*., 2018). Thus, we cannot expect a rescue from heat stress due to infection by the *Wolbachia* strain used here in *D. nigrosparsa*. We noted that the absolute value of knockdown depends on ramping speed and that it has been a topic of debate what ramping speed to use (Santos *et al*., 2011), but that in the frame of this study not absolute knockdown but the performance of infected flies relative to that of uninfected and cured flies was important.

Thermal tolerance is one of many aspects in thermal biology. Another aspect is thermal preference. *Drosophila melanogaster* infected with *w*Mel preferred one-degree cooler temperature than uninfected flies and about one to four degrees cooler when infected with *w*MelPop or *w*MelCS (Truitt *et al*., 2018; Arnold *et al*., 2019). In uninfected *D. nigrosparsa*, the preferred temperature was at around 10 °C for laboratory-reared flies and up to 35 °C for field-captured flies (Tratter Kinzner *et al*., 2019). If *Wolbachia* infect this fly species, it might prefer lower temperatures like in infected *D. melanogaster*, which could reduce the prospect of *Wolbachia*-infected *D. nigrosparsa* in the face of increasing temperature.

The increased locomotion in *D. nigrosparsa* observed in larvae and in adults may help the host to quickly react to climate change by easing the move to other areas, but, on the other hand, it may increase the visibility for predators and energy loss. Increases in host’s activities have been reported also from other *Wolbachia* strains. Beetles *Callosobruchus chinensis* infected with *Wolbachia w*BruCon and *w*BruOri walked significantly more than uninfected ones, which might help increasing their chance for mating (Okayama *et al*., 2016). Mosquitoes *Aedes aegypti* infected with *w*MelPop had up to 2.5-fold increase in activity compared with uninfected ones (Evans *et al*., 2009).

We found significant differences in wing size and shape of *D. nigrosparsa* between infected and cured lines, but these differences were more likely due to genetic drift and not due to *Wolbachia* as the cured lines were subpopulations of infected lines and had been separated from their parent populations for five generations before the wing measurement. In addition, if *Wolbachia* affect wing morphology, we would observe similar changes in those cured lines once *Wolbachia* were removed. Effects of genetic drift in *Drosophila* can occur within a few generations, for example, in *Drosophila subobscura* (Santos *et al*., 2013).

Our study indicated that *D. nigrosparsa* could be a host for *Wolbachia* like *Drosophila melanogaster,* the native host of *Wolbachia w*Mel, because vertical transmission is possible in this species. On the long term, the transmission of *Wolbachia* in *D. melanogaster* may be better than in *D. nigrosparsa* because *D. melanogaster* has a higher oviposition rate and a better tolerance of warm temperatures than *D. nigrosparsa* (Kinzner *et al*., 2018), both of which could increase the chance for horizontal transfer. This is because horizontal transfer is a stochastic event, and an infected host is therefore more likely to transfer *Wolbachia* to a new host species if there are more infected hosts available and if the number of *Wolbachia* cells is higher per host.

Here, we report effects of *Wolbachia w*Mel on *D. nigrosparsa* as a novel host. We observed low *Wolbachia* titer, possible cytoplasmic incompatibility, and increased locomotion in both larvae and adults. *Drosophila nigrosparsa* will suffer from an increasing temperature independently of whether uninfected (Kinzner *et al*., 2019) or infected, as *Wolbachia* had no impact on heat tolerance (this paper). However, *Wolbachia w*Mel might provide some benefits to this fly such as resistance to viruses or nutrition supplements, which both will be interesting to analyze in the future. In addition, some additional experiments such as longevity should be tested in the future. Finally, infection by *Wolbachia* strains other than *w*Mel may trigger different effects in this alpine vinegar fly.

## Acknowledgments

We thank Yuk-Sang Chan for teaching and partly performing the microinjection technique, Julien Martinez and Luis Teixeira for providing *Wolbachia* strains, and Peter Ladurner for providing a capillary puller; Marlene Haider, Martina Nindl, Julia Millinger, Leo Drechsel, Patrick Schwenter, and Ivan Ploner for helping with experiments; Magdalena Tratter Kinzner, Anja Ekblad, Philipp Andesner, and David Reiter for help with fly maintenance and laboratory work; Veronika Hierlmeier and Clemens P. Maylandt for comments on an earlier version of the manuscript; the University of Innsbruck for providing a doctoral grant to MD.

## Author Contributions

All authors conceived the ideas and design the study. MD performed the experiments, collected the data and performed the statistical analyses. MD, BCSS, and FMS wrote the manuscript. WA and FMJ revised and approved the manuscript.

## Data Accessibility

All data used in this manuscript are presented in the text and supporting information.

## Supplementary Data

**Table S1.** Cytoplasmic incompatibility and fecundity of *Drosophila nigrosparsa*.

**Table S2.** F-test and *t*-test between infected and cured lines in CTmax and CTmin experiments.

**Table S3.** Procrustes analysis of variance comparisons of centroid size between infected and uninfected lines.

**Table S4.** Procrustes analysis of variance comparisons of shape between infected and uninfected lines.

**Table S5.** Classification/misclassification table of discriminant analysis between infected and corresponding cured lines.

**Table S6.** Pearson correlation between mean centroid size and FA1 and FA2 indices for size fluctuation asymmetry between left and right wings.

**Table S7.** Shape asymmetry between left and right wings calculated from Procrustes analysis of variance.

